# Exploring the Complexity of Protein-Level Dosage Compensation that Fine-Tunes Stoichiometry of Multiprotein Complexes

**DOI:** 10.1101/700963

**Authors:** Koji Ishikawa, Akari Ishihara, Hisao Moriya

**Affiliations:** Research Core for Interdisciplinary Sciences, Okayama University, Okayama 700-8530, Japan; Course of Agrochemical Bioscience, Faculty of Agriculture, Okayama University, Okayama 700-8530, Japan

## Abstract

Proper control of gene expression levels upon various perturbations is a fundamental aspect of cellular robustness. Protein-level dosage compensation is one mechanism buffering perturbations to stoichiometry of multiprotein complexes through accelerated proteolysis of unassembled subunits. Although N-terminal acetylation- and ubiquitin-mediated proteasomal degradation by the Ac/N-end rule pathway enables selective compensation of excess subunits, it is unclear how dominant this pathway contributes to stoichiometry control. Here we report that dosage compensation depends only partially on the Ac/N-end rule pathway. Our analysis of genetic interactions between 18 subunits and 10 quality control factors in budding yeast demonstrated that multiple E3 ubiquitin ligases and N-acetyltransferases are involved in dosage compensation. We find that N-acetyltransferases-mediated compensation is not simply predictable from N-terminal sequence despite their sequence specificity for N-acetylation. We also find that the compensation of Pop3 and Bet4 is due in large part to a minor N-acetyltransferase NatD. Furthermore, canonical NatD substrates histone H2A/H4 were compensated even in its absence, suggesting N-acetylation-independent compensation. Our study reveals the complexity and robustness of the stoichiometry control system.

## Author Summary

Quality control of multiprotein complexes is important for maintaining homeostasis in cellular systems that are based on functional complexes. Proper stoichiometry of multiprotein complexes is achieved by the balance between protein synthesis and degradation. Recent studies showed that translation efficiency tends to scale with stoichiometry of their subunits. On the other hand, although protein N-terminal acetylation- and ubiquitin-mediated proteolysis pathway is involved in selective degradation of excess subunits, it is unclear how dominant this pathway contributes to stoichiometry control due to the lack of a systematic investigation using endogenous proteins. To better understand the landscape of the stoichiometry control system, we examined genetic interactions between 18 subunits and 10 quality control factors (E3 ubiquitin ligases and N-acetyltransferases), in total 100 combinations. Our data suggest that N-acetyltransferases are partially responsible for stoichiometry control and that N-acetylation-independent pathway is also involved in selective degradation of excess subunits. Therefore, this study reveals the complexity and robustness of the stoichiometry control system. Further dissection of this complexity will help to understand the mechanisms buffering gene expression perturbations and shaping proteome stoichiometry.

## Introduction

Controlling intracellular protein concentration at the proper level is a critical aspect of cellular systems. For example, it has been suggested that the cells regulate proteome concentration for a constant rate of biochemical reactions [1–3]. Moreover, since a wide variety of stress response pathways dynamically change the expression of proteins responsible for survival in various challenging environments, it is obvious that the control of protein concentration is also important for coping with perturbations and maintaining cellular homeostasis.

Genome-wide studies measuring cellular robustness against genetic perturbations showed that *Saccharomyces cerevisiae* cells are fragile against protein overexpression of a subset of the genome [4, 5]. This finding indicates that the cell system is generally robust against genetic perturbations to various biological processes. However, it is unclear how cells buffer such environmental changes [6]. We recently reported that upon an increase in gene copy number, approximately 10% of the yeast genome are transcribed linearly into mRNA levels but not directly translated into protein levels [7]. Comprehensive analyses of aneuploid yeast and mammalian cells also showed that approximately 20% of the genome exhibit this molecular phenotype [8, 9]. Therefore, this phenomenon known as protein-level dosage compensation may partially explain the buffering of gene expression perturbations.

Protein-level dosage compensation predominantly targets genes encoding subunits of multiprotein complexes [7–10]. The concentration of the dosage-compensated proteins bidirectionally changes in response to that of its partner subunits [7, 11]. These observations are consistent with a hypothesis predicting the deleterious effects due to stoichiometric imbalance of the complex subunits on cell growth [5, 6, 12]. Another line of evidence for this prediction comes from ribosome profiling analyses revealing a proportional synthesis strategy that enables translation efficiency to scale with subunit stoichiometry [13, 14]. This finding shows that stoichiometric balance of multiprotein complex subunits is mainly maintained at the translational level. In addition, this strategy is conserved from bacteria to higher eukaryotes [14]. In this context, dosage compensation can be recognized as a fail-safe mechanism that fine-tunes proteome stoichiometry [15].

The mechanism of dosage compensation is accelerated degradation of unassembled subunits by the ubiquitin–proteasome system [7, 11, 16, 17]. Multiple E3 ubiquitin ligases involved in stoichiometry control were previously identified [11, 18, 19], but we lack an understanding of their relative contribution to the compensation of identical subunits [20]. Furthermore, the underlying mechanism for selective degradation of excess subunits is not well understood. Although previous studies found evidence for selective compensation of Cog1 and Hcn1 in yeast and PLIN2 in mammalian cells by the Ac/N-end rule pathway [11, 21], it is unclear how dominant this pathway contributes to the compensation. The Ac/N-end rule pathway is one mechanism for protein degradation through the acetylation of N-terminal amino acid by N-acetyltransferases (NATs). With the limited number of examples, N-acetylation was shown to act as a degradation signal (N-degron) recognized by specific E3 ubiquitin ligases (N-recognin) [11, 18, 22]. It has been hypothesized that N-degron of unassembled subunit is exposed and recognized by N-recognin, whereas that of assembled subunit is shielded by the other subunits in the same complex and is inaccessible by N-recognin (Fig 1A) [11, 18, 23]. This model is consistent with the biological role of dosage compensation, but it remains to be investigated whether the compensation depends on this pathway.

**Fig 1.**
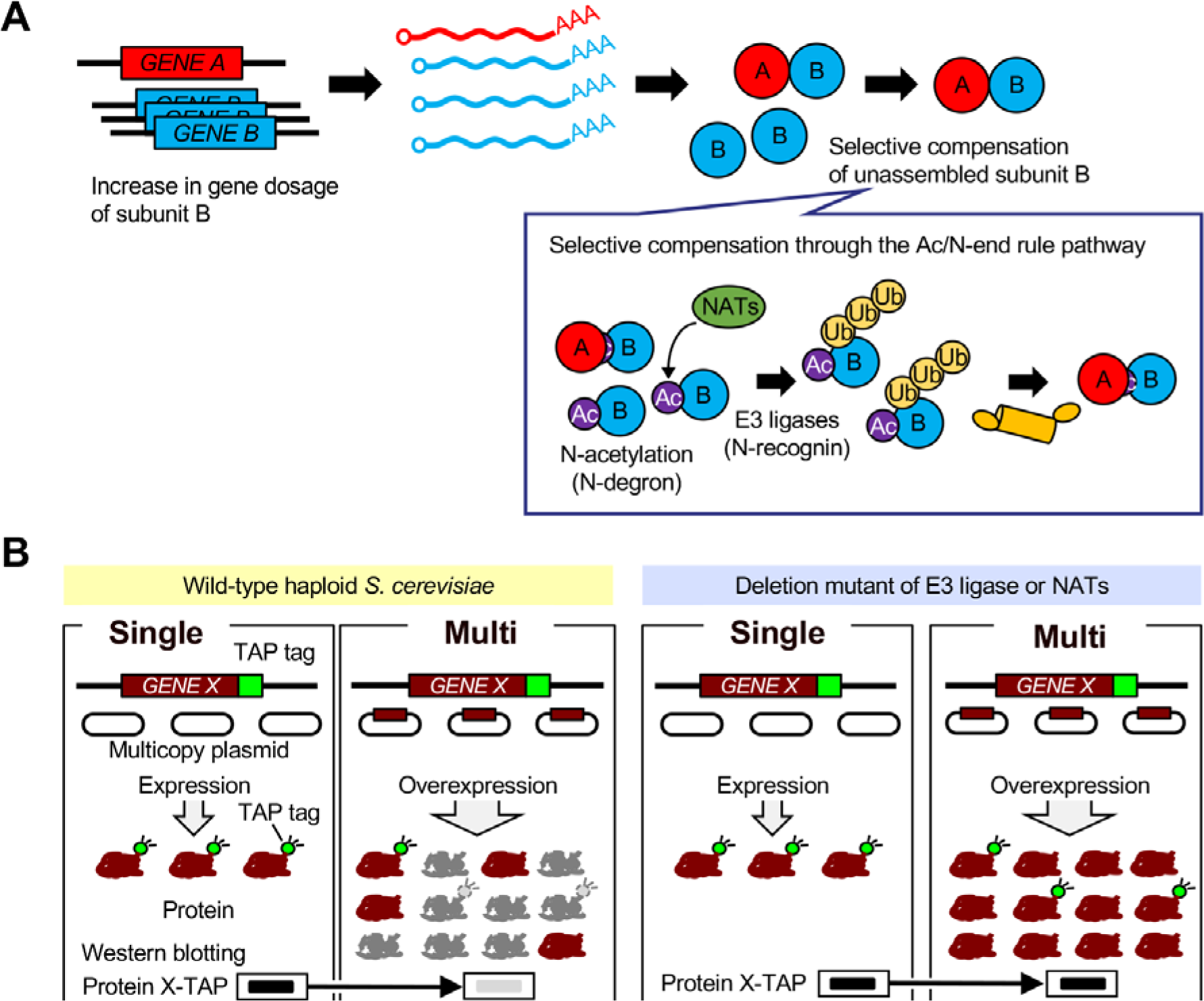
Genetic perturbation analysis of the stoichiometry control system. (**A**) Protein-level dosage compensation predominantly targets subunits of multiprotein complexes. The Ac/N-end rule pathway is involved in the underlying mechanism for the selective compensation of unassembled subunits. (**B**) An experimental setup for measuring the contribution of each quality control factor to dosage compensation. The untagged dosage-compensated protein was expressed from multicopy plasmid pTOW40836 containing the native regulatory sequences, including promoter and 5’ and 3’ untranslated regions. If the target gene is subjected to dosage compensation, the level of the TAP-tagged target protein expressed only from the chromosome decreases upon an increase in gene copy number (Multi) compared to vector control (Single). The Multi/Single ratio is higher than that in WT cells in the deletion mutant of each quality control factor (gene encoding E3 ubiquitin ligase or a catalytic subunit of NATs) if the deleted factor contributes to the compensation of the target gene.

In this study, we measured the contribution of E3 ubiquitin ligases and NATs to dosage compensation of the RNase P/MRP subunits, which are the compensated proteins identified in our previous study [7], in a systematic manner. We find that multiple E3 ubiquitin ligases and NATs are involved in the compensation. We also find that the dependency of the compensation on NATs is variable among tested subunits. Our findings suggest that dosage compensation does not completely depend on the Ac/N-end rule pathway and highlight the complexity of the stoichiometry control system.

## Results

### Multiple E3 ubiquitin ligases Tom1 and Not4 are involved in dosage compensation of identical subunits

To investigate which E3 ubiquitin ligases are involved in dosage compensation and their relative contribution, we measured protein levels of five dosage-compensated genes (*RBG1*, *MTW1*, *POP5*, *SAW1*, and *ERP2*), which were identified in our previous study [7], in five deletion mutants of E3 genes (*DOA10*, *TOM1*, *UBR1*, *SAN1*, and *NOT4*) involved in the protein quality control system including the N-end rule pathways [11, 18, 19, 24, 25]. Using a previously developed method for identifying genes with dosage compensation (S1A Fig) [7], we found that the amount of Pop5 in *tom1*∆ and *not4*∆ and Saw1 in *doa10*∆ was increased compared to wild type (WT) cells (S1B and S1C Fig).

Pop5 is a subunit of the RNase P/MRP complexes comprising of eleven subunits [26]. These complexes share eight subunits: Pop1, Pop3, Pop4, Pop5, Pop6, Pop7, Pop8, and Rpp1, while Rpr2 is included only in the RNase P and Snm1 and Rmp1 are included only in the RNase MRP. Most subunits of these complexes are subjected to dosage compensation [7], which prompted us to examine whether Tom1 and Not4 are involved in the compensation of the other subunits. We used an experimental setup by which the TAP-tagged target protein expressed from only one genomic locus is detected upon an increase in copy number of untagged version of the same gene (Fig 1B). We first compared protein levels between with and without overexpression in WT, *tom1*∆, or *not4*∆ cells (S2A Fig). This analysis showed a significant decrease in the compensation of Pop4, Pop5, Pop6, Pop7, Pop8, and Rpp1 in *tom1*∆ cells (Fig 2A and 2B), as well as Pop4, Pop5, Pop7, Rpp1, and Rmp1 in *not4*∆ cells (Fig 2D and 2E). In addition, the lower compensation of Rpr2 and Rmp1 in *tom1*∆ cells were marginally significant (*p*<0.1, two-tailed Welch’s *t* test) (S3 Fig). Therefore, Tom1 and Not4 were involved in dosage compensation of the RNase P/MRP subunits: Tom1 and Not4 target at least 6 and 5 out of 11 subunits, respectively (Table 1). While the sum of the contribution of these E3 ligases to the compensation of Pop7, Pop8, and Rpp1 reached to uncompensated protein level (Fig 2G), that of the other subunits did not reach to this level, which suggests the existence of unidentified E3 ligase(s) responsible for stoichiometry control.

**Table 1.** Responsible E3 ubiquitin ligases and NATs for dosage compensation of the RNase P/MRP subunits.

| Subunit | N-terminal first five amino acids | Responsible E3 Tom1 or Not4 | Responsible NATs Sequence-based | Responsible NATs Experiment-based |
| --- | --- | --- | --- | --- |
| Pop1 | MSGSL | n.d. | NatA | n.d. |
| Pop3 | MSGSL | not identified | NatA | NatD |
| Pop4 | MDRTQ | Tom1, Not4 | NatB | NatA |
| Pop5 | MVRLK | Tom1, Not4 | NatA | NatA<sup>b</sup> |
| Pop6 | MINGV | Tom1 | NatC | NatB |
| Pop7 | MALKK | Tom1, Not4 | NatA | not identified |
| Pop8 | MGKKT | Tom1 | NatA | NatB |
| Rpp1 | MLVDL | Tom1, Not4 | NatC | not identified |
| Rpr2 | MGKKA | Tom1<sup>a</sup> | NatA | NatB |
| Snm1 | MNKDQ | n.d. | NatB | n.d. |
| Rmp1 | MDEMD | Tom1<sup>a</sup>, Not4 | NatB | NatA, NatB |

<sup>a</sup> Marginally significant ( $p < 0.1$ , two-tailed Welch's $t$ test)
<sup>b</sup> Pop5 was significantly more compensated in *naa10Δ* than in WT cells.
n.d.: not detected

**Fig 2.**
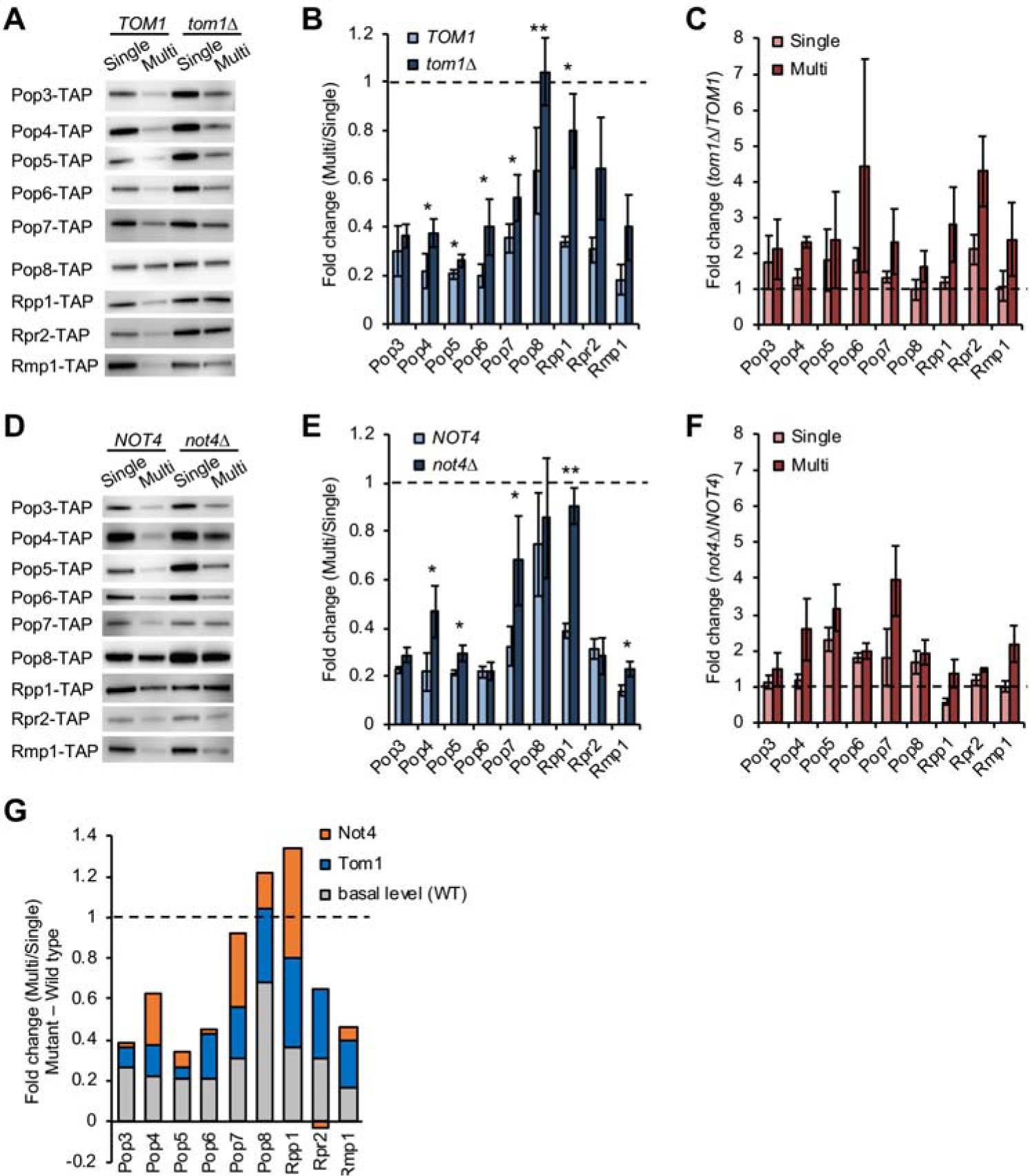
Tom1 and Not4 are involved in dosage compensation. (**A**) Western blots of the RNase P/MRP subunits in *tom1*∆ cells. The TAP-tagged subunit expressed from the chromosome was detected with PAP. Pop1 and Snm1 were not detected in our experiments. A representative blot from three biological replicates is shown. (**B**) Quantification of protein levels in the Multi condition relative to those in the Single condition in WT and *tom1*∆ cells. The average fold change ± s.d. was calculated from three biological replicates. Statistical significance was determined by a two-tailed Welch’s *t* test (\**p*<0.05, \*\**p*<0.01). Dashed line represents the same expression level between the Single and Multi conditions. (**C**) Comparison of protein levels between WT and *tom1*∆ cells in the Single or Multi conditions. The average fold change ± s.d. was calculated from three biological replicates. Dashed line represents the same expression level between WT and *tom1*∆ cells. (**D–F**) Same as in (**A–C**), except that shown are Western blots of the RNase P/MRP subunits in *not4*∆ cells (**D**) and their quantification (**E, F**). (**G**) The relative contribution of Tom1 and Not4 to the compensation. The basal level is the average fold change in WT cells. The contribution of Tom1 and Not4 was calculated as the average fold change in each mutant minus the basal level.

We next compared protein levels between WT and each mutant with or without overexpression (S2B Fig). This analysis showed the higher levels of Pop6 and Rpr2 in *tom1*∆ and Pop5 and Pop6 in *not4*∆ than those in WT cells without an increase in gene copy number (*p*<0.05, two-tailed Welch’s *t* test) (Fig 2C and 2F, Single). As this comparison is between vector controls in WT and mutant cells, Tom1 and Not4 may be involved in degradation of these subunits in unperturbed conditions. In addition, since Pop6 was not subjected to Not4-mediated compensation, Not4 contributes to its degradation but this is not accelerated upon overexpression.

### Dosage compensation of Pop6 and Pop7 occurs in response to their intracellular concentration

Since both Pop6 and Pop7, which form the Pop6–Pop7 heterodimeric subcomplex [27, 28], are subjected to dosage compensation (Fig 2), the amount of Pop6 and Pop7 may change in response to that of partner subunits. We hypothesized that high *POP7* dosage leads to changes in the fraction of the stable/assembled and unstable/unassembled pools of Pop6 (Fig 3A, top). Furthermore, since the mechanism of dosage compensation is accelerated degradation of unassembled subunits, the balance of these subunit pools may be perturbed in the absence of Tom1 or Not4 (Fig 3B, top). As expected (Fig 3A and 3B, bottom), we found that Pop6 was stabilized by an increase in *POP7* dosage, and vice versa, in WT, *tom1*∆, or *not4*∆ cells (Fig 3C and 3D). These results indicate that dosage compensation of Pop6 and Pop7 occurs bidirectionally in response to changes in intracellular concentration of each subunit: accelerated degradation of Pop6/Pop7 upon high *POP6*/*POP7* dosage (downward compensation) and reduced degradation of Pop6/Pop7 upon high *POP7*/*POP6* dosage (upward compensation).

**Fig 3.**
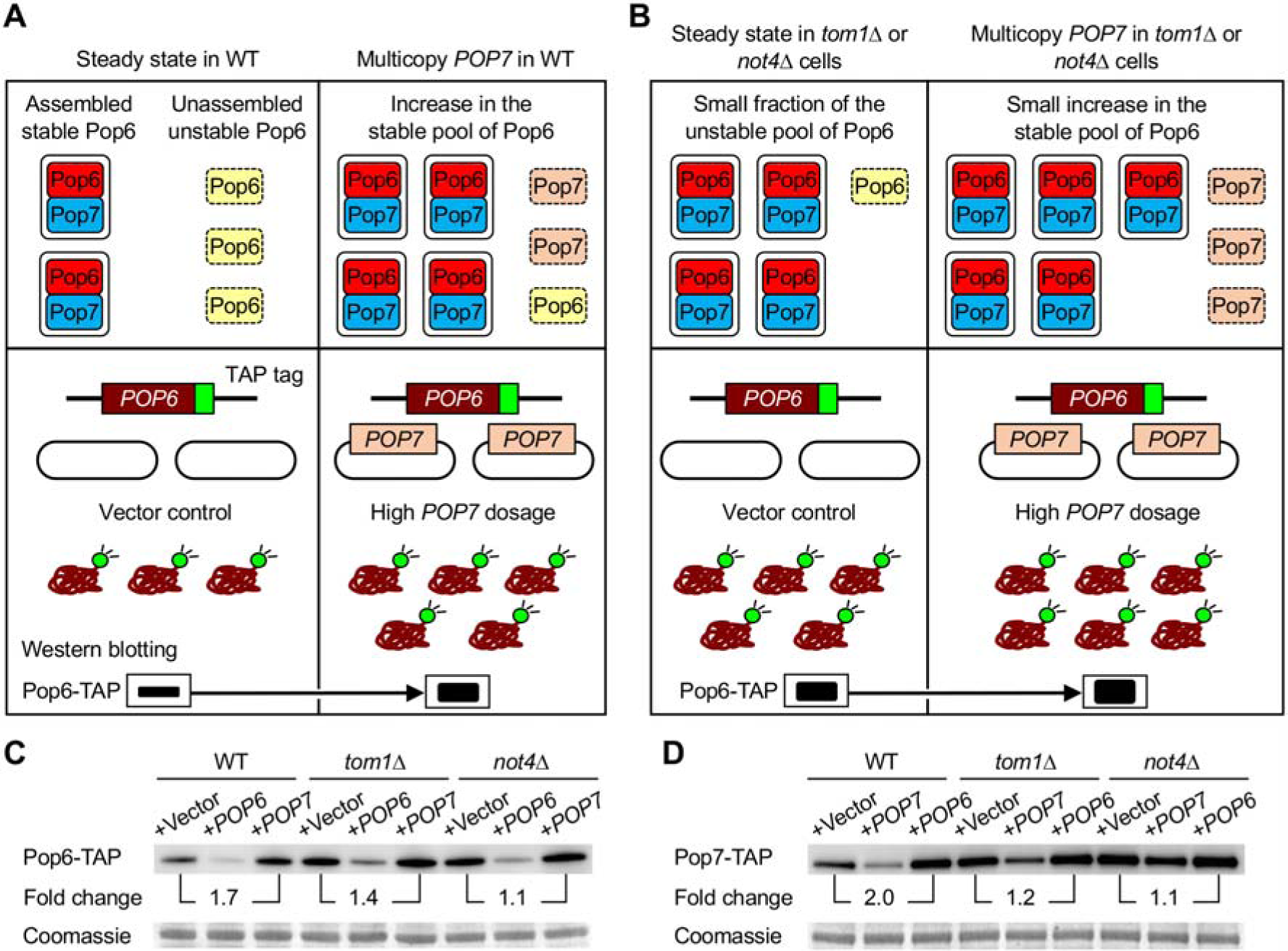
Bidirectional dosage compensation of the Pop6–Pop7 heterodimeric subcomplex. (**A**) A model for stoichiometry control of Pop6 and Pop7 in WT cells (top) and an experimental setup for testing this model (bottom). Pop6 becomes more stable when in complex with the partner subunit Pop7, while unassembled Pop6 is less stable (top left). Upon multicopy expression of Pop7, potentially degraded Pop6 becomes stable by forming the complex with excess Pop7 (top right). Pop6 level in vector control is the steady state level (bottom left). If Pop6 turnover is reduced by high *POP7* dosage, Pop6 level is increased compared to that in vector control (bottom right). (**B**) Same as in (**A**), except that shown is the case of *tom1*∆ or *not4*∆ cells. Potentially degraded Pop6 in WT cells becomes stable in the absence of Tom1 or Not4 responsible for Pop6 and Pop7 degradation, leading to an increase in the Pop6–Pop7 complex (top left). Upon multicopy expression of Pop7, the remaining unstable pool of Pop6 becomes stable by forming the complex with excess Pop7 (top right). The steady state level of Pop6 in *tom1*∆ or *not4*∆ cells is higher than that in WT cells (bottom left). If the turnover of Pop6 is further reduced by high *POP7* dosage, it leads to a small increase in Pop6 level (bottom right). (**C, D**) Western blots of Pop6-TAP (**C**) and Pop7-TAP (**D**) in WT, *tom1*∆, and *not4*∆ cells. The TAP-tagged proteins were detected with PAP in the Single (+Vector: pTOW40836) and Multi (+*POP6*: pTOW40836-*POP6*, +*POP7*: pTOW40836-*POP7*) conditions. Quantification of band intensities relative to the Single condition in each strain is shown. Coomassie staining of a 50-kDa protein, corresponding to enolase, is shown as a loading control.

Upon increasing gene copy number of the partner subunit, the relative ratio of Pop6 and Pop7 in *tom1*∆ and *not4*∆ was lower than that in WT cells. This difference may reflect the regulation of the compensation degree, as the amount of Pop6 and Pop7 was increased in both mutants without changes in gene copy number (Fig 3C and 3D, +Vector). Therefore, Tom1 and Not4 impact the balance of stable/unstable subunit pools of the Pop6–Pop7 heterodimeric subcomplex.

### NATs contribute to dosage compensation in a complex manner

*S. cerevisiae* has five NATs: NatA, NatB, NatC, NatD, and NatE whose catalytic subunits are respectively Naa10, Naa20, Naa30, Naa40, and Naa50 [29]. NATs have different sequence specificities for N-acetylation and their substrates can be classified by the first two N-terminal residues except NatD that recognizes approximately five-residue sequence motifs [18, 29, 30]. As shown in Fig 4A, based on a prediction from N-terminal amino acids, the RNase P/MRP subunits can be N-acetylated by NatA, NatB, or NatC. We examined whether the RNase P/MRP subunits are less compensated in *naa10*∆, *naa20*∆, *naa30*∆, *naa40*∆, or *naa50*∆ cells using the same experimental setup described in Fig 1B. Our experiments showed that Pop4 and Rmp1 in *naa10*∆ (Fig 4B and 4C) and Pop6, Pop8, Rpr2, and Rmp1 in *naa20*∆ (Fig 4D and 4E) were significantly less compensated than those in WT cells, while all tested subunits were compensated in *naa30*∆ or *naa50*∆ as well as in WT cells (Fig 4F, 4G, 4J, and 4K). Only Pop3 was uncompensated in *naa40*∆ cells (Fig 4H and 4I), and the half-life of Pop3 was prolonged upon increasing *POP3* copy number in *naa40*∆ cells (S4A and S4B Fig). Of note, Pop3 was stabilized in *naa10*∆ compared to WT cells with or without an increase in gene copy number (S5 Fig) (*p*<0.05, two-tailed Welch’s *t* test), suggesting the involvement of NatA in Pop3 degradation even in unperturbed conditions. We also found that Pop5 compensation was more pronounced in *naa10*∆ than in WT cells (Fig 4B and 4C), suggesting that accelerated Pop5 degradation during dosage compensation is blocked by NatA.

**Fig 4.**
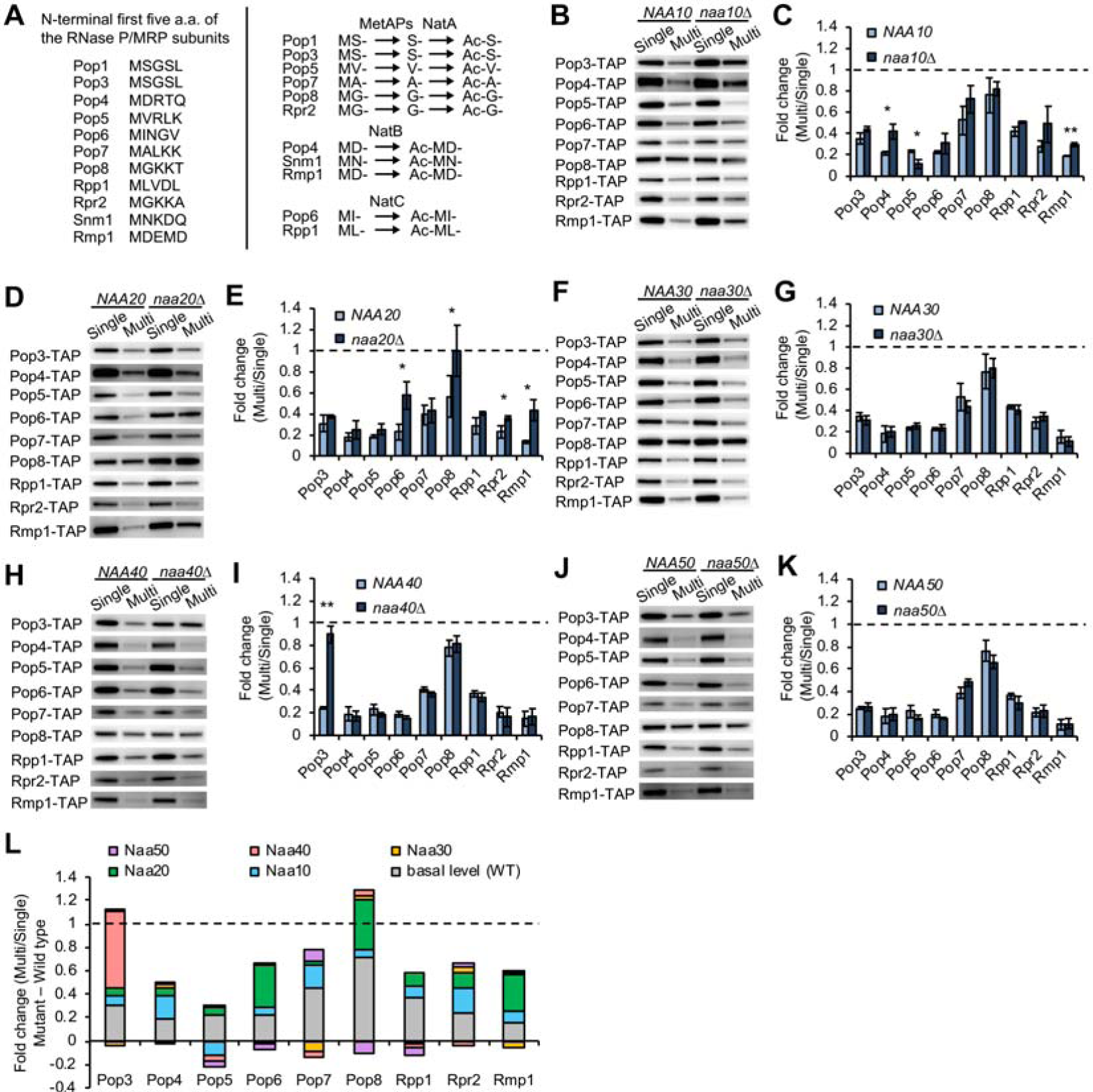
NATs contribute to dosage compensation in a complex manner. (**A**) The first five amino acids of the RNase P/MRP subunits (left). Based on these sequences, NatA, NatB, or NatC are responsible for their N-acetylation (right). NatA-mediated acetylation requires removal of N-terminal Methionine by Met-aminopeptidases Map1 and Map2 (MetAPs). (**B**) Western blots of the RNase P/MRP subunits in *naa10*∆ cells. The TAP-tagged subunit expressed from the chromosome was detected with PAP. Pop1 and Snm1 were not detected in our experiments. A representative blot from three biological replicates is shown. (**C**) Quantification of protein levels in the Multi condition relative to those in the Single condition in WT and *naa10*∆ cells. The average fold change ± s.d. was calculated from three biological replicates. Statistical significance was determined by a two-tailed Welch’s *t* test (\**p*<0.05, \*\**p*<0.01). Dashed line represents the same expression level between the Single and Multi conditions. (**D–K**) Same as in (**B, C**), except that shown are Western blots of the RNase P/MRP subunits and their quantification in *naa20*∆ (**D, E**), *naa30*∆ (**F, G**), *naa40*∆ (**H, I**), or *naa50*∆ (**J, K**) cells. (**L**) Summary of the contribution of NATs to dosage compensation of the RNase P/MRP subunits. The basal level is the average fold change in WT cells. The contribution of each NAT was calculated as the average fold change in each mutant minus the basal level.

We summarized the contribution of each NAT, leading to several findings (Fig 4L; Table 1). First, NatB-mediated compensation of Rmp1 is the only case that can be expected from its N-terminal sequence. Second, conversely, the compensation of Pop4 and Pop6 were engaged with NatA and NatB but not NatB and NatC, respectively. Additionally, Pop8 and Rpr2 were subjected to the compensation mediated by NatB but not NatA. Third, Rmp1 was compensated through multiple NATs, NatA and NatB. Fourth, the compensation of Pop3 was due in large part to NatD but not NatA. Fifth, NATs were not identified for the compensation of Pop7 and Rpp1. Finally, NATs-mediated compensation of the RNase P/MRP subunits was mainly mediated by NatA and NatB, although their impact on protein levels was limited. Taken together, the N-terminal sequence cannot simply explain NATs-mediated dosage compensation. In all cases except for Pop3 and Pop8, the sum of the contribution of each NAT did not reach to uncompensated protein level, suggesting that the stoichiometry control system depends only partially on NATs-mediated dosage compensation.

### The first two N-terminal residues are not sufficient to determine the combination of NATs and E3 ligases in the Ac/N-end rule pathway

By summing the contribution of the tested E3 ligases and NATs to dosage compensation of the RNase P/MRP subunits, the protein level of all subunits except Pop5 reached to the uncompensated level (Fig 5A). Pop5 seems to be compensated through NATs-independent degradation pathway predominantly mediated by redundant control via Tom1 and No4 and/or unidentified E3 ligase(s). Pop4 and Pop7 were less compensated to the same degree in both *not4*∆ and *naa10*∆ cells (Fig 5). Because Not4 was previously identified as an N-recognin [11], these subunits may be compensated by the combination of NatA and Not4. Both Pop4 and Rmp1 are MD-starting proteins, but Rmp1 was compensated by not only NatA and Not4 but also NatB (Fig 2 and 4). Cog1 is also a MD-starting protein whose protein level is compensated by NatB and Not4 upon its overexpression [11]. These observations suggest that even though the first two N-terminal residues of substrate proteins are the same, the compensation through the Ac/N-end rule pathway occurs by different combinations of NATs and E3 ligases. Moreover, Rpp1 was almost fully uncompensated in *not4*∆ but not in all NATs mutants (Fig 2D and 2E), indicating that Not4-mediated compensation does not necessarily require NATs.

**Fig 5.**
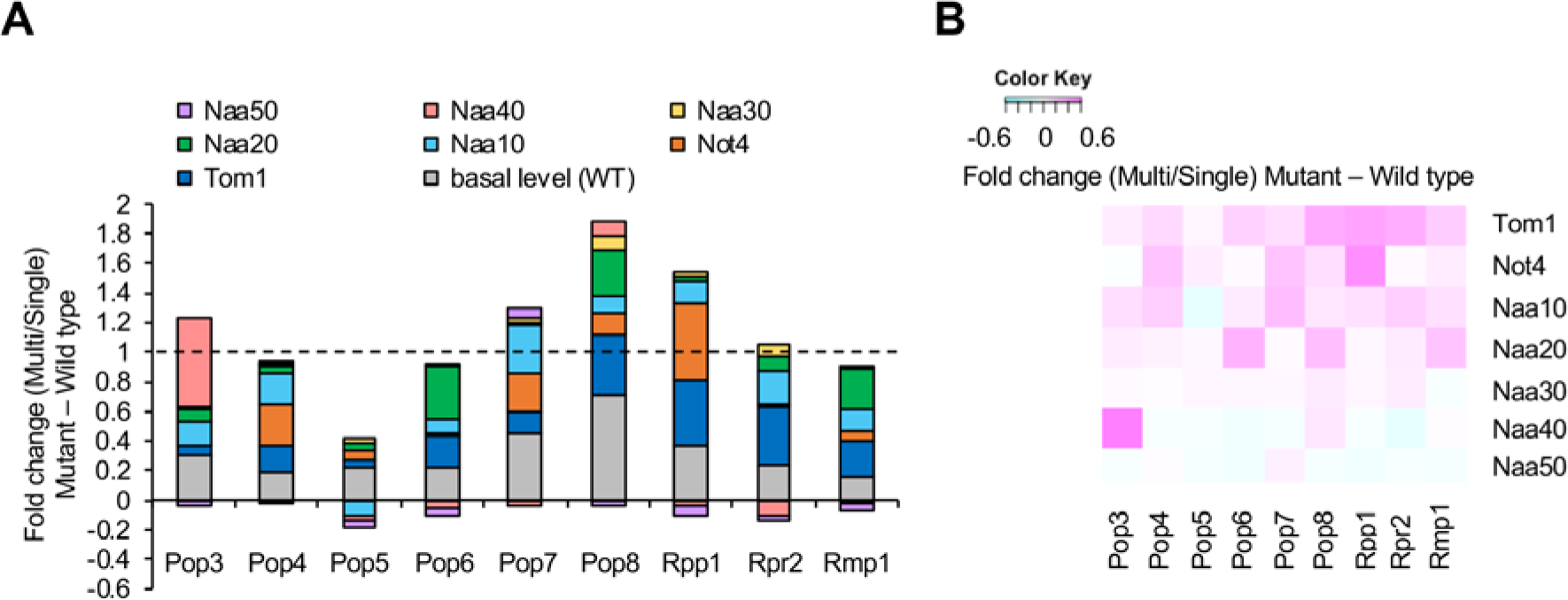
Relative contribution of Tom1, Not4, and NATs to dosage compensation of the RNase P/MRP subunits. (**A, B**) Summary of the contribution of Tom1, Not4, and NATs to dosage compensation of the RNase P/MRP subunits. The basal level is the average fold change in WT cells. The contribution of each factor was calculated as the average fold change in each mutant minus the basal level. Bar graph and the heat map show the same values.

### Dosage compensation of histone H2A/H4 may not depend on the Ac/N-end rule pathway

As shown in Fig 4H and 4I, NatD efficiently contributes to the compensation of Pop3 that is not identified as a NatD substrate [31]. To further examine whether NATs-mediated dosage compensation is hardly predictable from N-terminal sequence, we next analyzed canonical NatD substrates. NatD is more selective than the other NATs [29], and only histone H2A/H4 subunits are identified as its substrates [31–33]. These histone subunits are MS-starting proteins that can be NatA substrates, but they are N-acetylated in the absence of Naa10 or its auxiliary subunit Naa15 [34]. Thus, we first tested whether NatD is involved in the compensation of histone H2A/H4 subunits: Hta2, Hhf1, and Hhf2. We found that they were subjected to dosage compensation even in the absence of Naa40 (Fig 6A–6C), suggesting that NatD is involved in their N-acetylation but not degradation. Additionally, the compensation was observed not only in *naa40*∆ but also in *naa10*∆, *naa20*∆, or *naa30*∆ as well as in WT cells (S6 Fig). We also found that they all were significantly less compensated in *tom1*∆ cells, although the degree of compensation was different among them (Fig 6A–6C). These results suggest that Tom1-mediated compensation of Hta2, Hhf1, and Hhf2 does not require NATs activity and that N-acetylation and dosage compensation are not necessarily linked. Therefore, dosage compensation of histone H2A/H4 may be mediated by N-acetylation-independent pathway.

**Fig 6.**
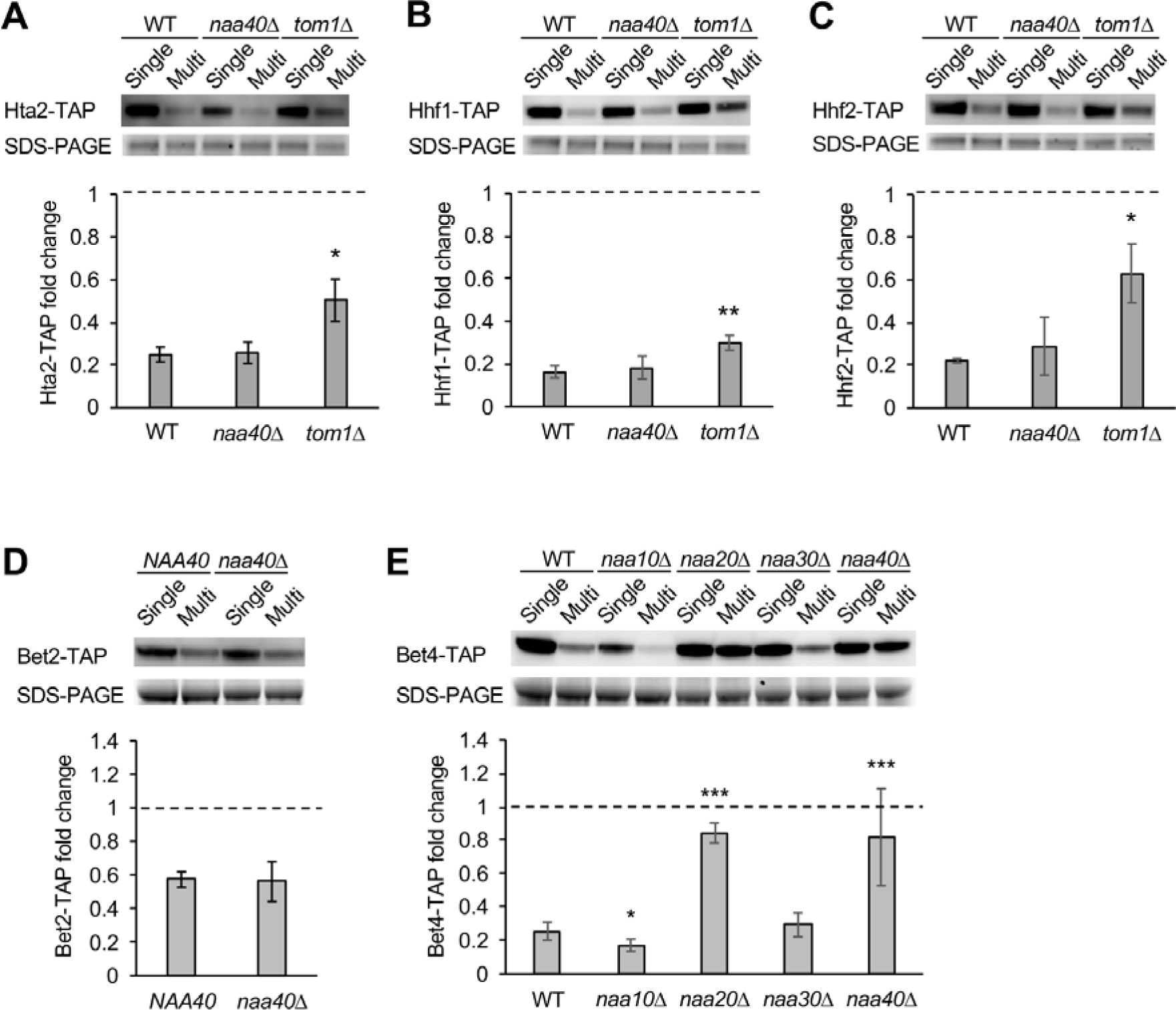
Dissection of the complexity of NATs-mediated dosage compensation by focusing on NatD. (**A–C**) Tom1-mediated dosage compensation of NatD substrates does not require Naa40. Western blot analysis of histone H2A/H4: Hta2-TAP (**A**), Hhf1-TAP (**B**), and Hhf2-TAP (**C**) in WT, *naa40*∆, or *tom1*∆ cells. The TAP-tagged subunits were detected with PAP (top). The average fold change ± s.d. was calculated from three biological replicates (bottom). Statistical significance was determined by a two-tailed Welch’s *t* test (\**p*<0.05, \*\**p*<0.01). Dashed line represents the same expression level between the Single and Multi conditions. (**D**) Western blot analysis of Bet2-TAP in *naa40*∆ cells. The TAP-tagged subunits were detected with PAP (top). The average fold change ± s.d. was calculated from four biological replicates (bottom). (**E**) Western blot analysis of Bet4-TAP in *naa10*∆, *naa20*∆, *naa30*∆, and *naa40*∆ cells. The TAP-tagged subunits were detected with PAP (top). The average fold change ± s.d. was calculated from eight and three biological replicates for *naa40*∆ and the other mutants, respectively (bottom) (\**p*<0.05, \*\*\**p*<0.001).

### NatD-mediated dosage compensation of a MH-starting protein Bet4

The selectivity of NatD-mediated N-acetylation comes from a sequence motif recognized by NatD and its substrate recognition site tailored for this motif [30]. The N-terminal of Pop3 is SGSL after removal of N-terminal Met by Met-aminopeptidases, which is different from the known motifs involved in histone subunits: SGGK (H2A) and SGRG (H4) [31]. We thus examined whether Bet2, which is a MSGSL-starting protein and a subunit of the Bet2–Bet4 heterodimer [35], is subjected to NatD-mediated compensation. We found that Bet2 was compensated by the same degree in either *naa40*∆ or WT cells (Fig 6D), indicating that the first MSGSL is not sufficient to trigger NatD-mediated compensation. Unexpectedly, Bet4 was remarkably less compensated in *naa40*∆ compared to WT cells (Fig 6E). Bet4 was indeed stable upon increasing *BET4* copy number in *naa40*∆ cells due to the prolonged half-life (S4C and S4D Fig). This phenomenon was surprising because Bet4 is a MH-starting protein that is virtually not N-acetylated while N-acetylation of Bet4 was detected but its responsible NATs were not identified [29, 36].

We further examined the contribution of the other NATs to Bet4 compensation and found that Bet4 was significantly less compensated in *naa20*∆ cells (Fig 6E). Thus, NatB and NatD are involved in Bet4 compensation. We also found an increase in Bet4 compensation in *naa10*∆ cells (Fig 6E), suggesting a NatA-mediated block of Bet4 degradation during the compensation. Although each of them is similar to Rmp1 in *naa10*∆ and *naa20*∆ and Pop5 in *naa10*∆ cells (Fig 4B–4E), both effects of NATs deletion were observed only for Bet4. Therefore, Bet4 was identified as the compensated protein whose stoichiometry is bidirectionally controlled by multiple NATs upon genetic perturbations.

## Discussion

To investigate how well the Ac/N-end rule pathway explains dosage compensation, we systematically measured the contribution of E3 ubiquitin ligases and NATs to the compensation of the RNase P/MRP subunits (Fig 2 and 4). Our data consisting of 18 subunits and 10 quality control factors, in total 100 combinations, demonstrated that multiple E3 ubiquitin ligases and NATs are involved in the compensation. NATs-mediated compensation was observed for 7 out of 14 subunits of the RNase P/MRP, histone, and Bet2–Bet4 complexes (Fig 4 and 6). However, given that the lack of NATs did not completely reduce the compensation of the tested subunits except for Pop3 in *naa40*∆ and Bet4 in *naa20*∆ and *naa40*∆ cells, stoichiometry control of multiprotein complexes depends only partially on the Ac/N-end rule pathway.

In this study, we manipulated gene copy number to perturb cellular systems and elucidate mechanisms for buffering gene expression perturbations. Our data showed that NATs contribute to proteolysis upon exogenous overexpression of the RNase P/MRP subunits, whereas without overexpression the endogenous protein levels were almost not affected in NATs mutants compared to E3 mutants (Fig 2C and 2F and S5 Fig). This is consistent with a recent comprehensive analysis showing that N-acetylation rarely acts as a degradation signal in physiological conditions in yeast [37]. In agreement with these findings, a limited number of subunits are physiologically synthesized in excess compared to stoichiometry and the half-life of such subunits tends to be faster than that of proportionally synthesized subunits [13, 14]. Therefore, NATs-mediated dosage compensation may be a fail-safe mechanism that is rarely and predominantly triggered by genetic perturbations or physiological overexpression causing stoichiometric imbalance.

We identified Tom1 as a factor of the stoichiometry control system (Fig 2A and 2B), which is consistent with Tom1-mediated degradation of excess ribosomal subunits [19]. It should be noted that, as discussed in [19], basic isoelectric point is a common characteristic of known Tom1 substrates (Cdc6, Hht2, Yra1, and Rpl26a) and the RNase P/MRP subunits except Pop8 (S7 Fig) [19, 38–40]. The weak electrical interaction between Tom1 with acidic isoelectric point and Pop8 might explain why the compensation degree of Pop8 is lower than the other subunits. Additionally, these substrates as well as Tom1 are nucleic acid binding and localize in the nucleus [41]. These characteristics suggest that Tom1 is broadly responsible for the compensation of nucleic acid binding subunits.

We found that ribosome-associated factors Not4 and NATs were involved in dosage compensation of the RNase P/MRP subunits (Fig 2D, 2E, and 4), which is consistent with the stabilization of Cog1 in *not4*∆ or *naa20*∆ cells upon its overexpression [11, 42–44]. These observations and cotranslational ubiquitination of a subset of the proteome suggest the possibility of cotranslational dosage compensation [45, 46]. This is also supported by the findings that multiprotein complex subunits tend to assemble cotranslationally and that such subunits are prone to aggregation and degradation in the absence of their partner subunits [47, 48]. Therefore, future investigations of this possibility may gain insight into the hierarchy of the multilayered stoichiometry control system.

We described the complexity of the stoichiometry control system (Fig 5 and 6, Table 1), as (i) multiple E3 ligases Tom1 and Not4 are involved in the compensation of Pop4, Pop5, Pop7, Rpp1, and Rmp1, (ii) N-terminal sequence is not necessarily a determinant of which NATs contribute to dosage compensation, and (iii) multiple NATs are involved in the compensation of Rmp1 and Bet4. Robust control of Rmp1 seems to be due to its C-terminal polylysine sequence CKKKKKRKKKNK that induces translation arrest coupled with Not4-mediated proteolysis [49]. Our findings thus suggest that the complexity of this system reflects its robustness. Furthermore, as argued in [50], other mechanisms for sensing unassembled subunits and correcting stoichiometric imbalance may exist. For example, the most recent study showed that protein aggregation of excess subunits functions as the compensation pathway in aneuploid yeast cells [51]. It is also possible to speculate that substrate recognition by one NAT is partially redundant with the other NAT because of the remarkable reduction of Bet4 compensation in both NatB and NatD mutants and functional redundancy between yeast NatC and human NatF (Fig 6E) [52].

The compensation of Pop3 in *naa40*∆ and Bet4 in *naa20*∆ and *naa40*∆ cells was almost fully reduced (Fig 4H, 4I, and 6E), supporting the dependency of stoichiometry control on the Ac/N-end rule pathway [11]. However, these are rare cases in our study and the first examples of NatD-mediated dosage compensation. Indeed, canonical NatD substrates histone H2A/H4 were compensated even in the absence of Naa40 (Fig 6A–6C). Importantly, consistent with the stabilization of histone H3 subunit Hht2 by *TOM1* deletion [39], H2A/H4 were less compensated in *tom1*∆ cells. These results suggest that unassembled subunits are selectively degraded by not only the Ac/N-end rule but also other pathways. This is further suggested by the observation that Rpp1 compensation involves Tom1 and Not4 but not NATs (Fig 5, Table 1).

We also found that Pop5 and Bet4 were destabilized when they were overexpressed in *naa10*∆ cells (Fig 4B, 4C, and 6E). Similarly, previous studies showed that proteins with unacetylated N-terminal Met followed by a small hydrophobic residue and MN-starting proteins are destabilized in *naa30*∆ and *naa20*∆ cells, respectively, by the Arg/N-end rule pathway [37, 53]. However, because N-terminal sequence of Pop5 (MV) and Bet4 (MH) are not appropriate for these cases, their degradation in *naa10*∆ cells might be accelerated by unknown mechanisms. We speculate that complex stoichiometry is robustly maintained by switching degradation pathways in response to perturbations in the stoichiometry control system. Indeed, the interplay between the Arg/N-end rule and the Ac/N-end rule pathways was proposed based on the observation that the short-lived reporter protein and Msn4 are synergistically stabilized in double mutants *naa20*∆ *ubr1*∆ and *naa30*∆ *ubr1*∆, respectively [53, 54].

Quality control of multiprotein complexes impacts a broad range of biological processes because of the fact that cellular systems are based on functional complexes. For example, the RNase P and MRP complexes are responsible for maturation of tRNA and rRNA, respectively [26]. Additionally, Pop1, Pop6, and Pop7 are shared with telomerase [55]. Our finding of robust control of dosage balance between Pop6 and Pop7 seems to be in line with their functional importance (Fig 3). More generally, dosage compensation may play a role as a fail-safe mechanism for shaping proteome stoichiometry as discussed above. In agreement with this concept, paralogous complex subunits tend to be compensated each other for modulating protein interactome [56, 57]. Furthermore, proteolysis-mediated compensation perversely corrects physiologically caused stoichiometric imbalance during meiosis [58], which may explain how cells cope with a subset of subunits overexpressed in unperturbed conditions [13]. Therefore, further dissection of the complexity and working principles of the stoichiometry control system will help to describe cellular robustness.

## Materials and Methods

### Strains, plasmids, and media

For construction of mutant strains lacking a gene encoding the E3 ubiquitin ligase (*TOM1* or *NOT4*) or N-aetyltransferase (*NAA10*, *NAA20*, *NAA30*, *NAA40*, or *NAA50*) and carrying a TAP-tagged gene of interest, the deletion collection and the TAP collection of haploid yeast (Dharmacon) were used. BY4741 carrying Pop4-TAP, Rmp1-TAP, or Bet4-TAP were constructed in this study. Genomic DNA of each single deletion strain was extracted, and then, each locus replaced with the *kanMX4* cassette was amplified by PCR with a primer library prepared in our previous study [5]. The TAP-tagged strains were transformed with the PCR products by the lithium acetate method and selected on YPD plates containing G418 (200 µg/mL). For construction of plasmids, DNA fragment of each target region was amplified from the genome and cloned into pTOW40836 by homologous recombination in BY4741 cells. The TAP-tagged strains described above were transformed with the plasmids and selected on SC medium lacking uracil.

### Measurement of plasmid copy number

The plasmid copy number was measured by the gTOW technique [59]. The cells grown at log-phase for Western blotting were harvested for the gTOW analysis from the same culture. In short, 200 µL of the culture was centrifuged and the pelleted cells were suspended in 100 µL of zymolyase solution [2.5 mg/mL Zymolyase-100T dissolved in 1.2 M sorbitol, 10 mM sodium phosphate (pH7.5)] and incubated at 37 °C for 15 min for DNA extraction. The suspension was heated at 100 °C for 10 min, cooled at –80 °C for 10 min, and again heated at 100 °C for 10 min. After cooling down to RT followed by a centrifugation, the supernatant was subjected to real-time quantitative PCR with Lightcycler 480 (Roche). The primers for *LEU3* on the genome or *leu2d* gene on pTOW40836 and SYBR Green I Master (Roche) were used for the PCR. The resulting plasmid copy number was calculated based on the expression levels of *LEU3* and *leu2d* genes according to the method described previously [59].

### Western blot analysis

Cells were grown at 30 °C in 2 mL of SC–Ura medium for overnight and then measured the optical density at 600 nm (OD_600_) and inoculated into fresh medium at initial OD_600_ of 0.5 in 3 mL. After 4 h, 1 OD_600_ units were harvested at log-phase when OD_600_ was around 1. The cells were treated with 1 mL of 0.2 N NaOH for 5 min at room temperature (RT) and centrifuged at 15,000 rpm for 1 min. The pelleted cells were suspended in 50 µL of 1× NuPAGE LDS Sample Buffer (Invitrogen) containing 100 mM DTT and heated at 100 °C for 5 min. For the analysis of the TAP-tagged proteins, two-fold serially diluted lysates were analyzed as described below, and we confirmed that 0.025 OD_600_ units provide appropriate signal in the linear range. The extract diluted 8-fold with 1× NuPAGE LDS Sample Buffer, corresponding with 0.025 OD_600_ units, was separated by polyacrylamide gel electrophoresis with lithium dodecyl sulfate (SDS-PAGE) on NuPAGE 4%–12% Bis-Tris Gel (Invitrogen). For the analysis of the GFP-tagged proteins, 0.2 OD_600_ units were subjected to SDS-PAGE. The separated proteins were transferred onto PVDF membrane using the iBlot Transfer Stack PVDF membrane (Invitrogen). The blotted membrane was treated with PBST [1× PBS, 0.1% Tween 20] for 10 min, and then blocked with 4% skim milk in PBST for 1 h at RT. The TAP-tagged proteins were probed with PAP (Sigma-Aldrich) (1:2,000) for 1 h at RT. The GFP-tagged proteins were probed with anti-GFP antibody (Roche) (1:1,000) and peroxidase-conjugated secondary antibody (Nichirei Biosciences) (1:1,000) for 1 h at RT. After washing the membrane with PBST for 5 min for three times, chemiluminescence was induced by adding 500 µL of SuperSignal West Femto Maximum Sensitivity Substrate (Thermo Scientific) on the membrane and detected using LAS-4000 image analyzer (Fujifilm) and ImageQuant LAS 4000 (GE Healthcare). Quantification of the band intensity was carried out using ImageQuant TL (GE Healthcare) and the fold change was calculated after background subtraction. After washing the membrane with sterile water for 5 min for three times, total proteins were visualized by Coomassie staining with SimplyBlue SafeStain (Invitrogen) to confirm equal loading of proteins. The stained membrane was digitized using LAS-4000 image analyzer and ImageQuant LAS 4000.

For the analysis of histone H2A/H4 and the Bet2–Bet4 heterodimer, the harvested cells were treated with 1 mL of 0.2 N NaOH for 5 min at RT. Cells were suspended in 1× NuPAGE LDS Sample Buffer (Invitrogen) and then heated at 70 °C for 10 min. The supernatant corresponding to 0.5 OD_600_ units was labeled with EzLabel FluoroNeo (ATTO) and subjected to SDS-PAGE, followed by Western blot with PAP (Sigma-Aldrich) (1:2,000) as described above.

### Cycloheximide chase assay

The degradation rate of Pop3 and Bet4 was measured as previously described [7]. Briefly, the log-phase culture corresponding to 1 OD_600_ unit of cells was harvested for time point 0 and afterwards CHX was added to a final concentration of 200 μg/mL. Cells were harvested after 1, 2, 4, and 6.5 h, followed by total protein extraction in 1× NuPAGE LDS Sample Buffer. The supernatant corresponding to 0.5 OD_600_ units was subjected to Western blot analysis using PAP as described above. The remaining protein level at each time point was calculated against that at time point 0.

## Acknowledgments

We thank the members of Moriya laboratory for their support.

