## Supplementary Material for "Exploring the Complexity of Protein-Level Dosage Compensation that Fine-Tunes Stoichiometry of Multiprotein Complexes"

**Supplementary Figure 1**

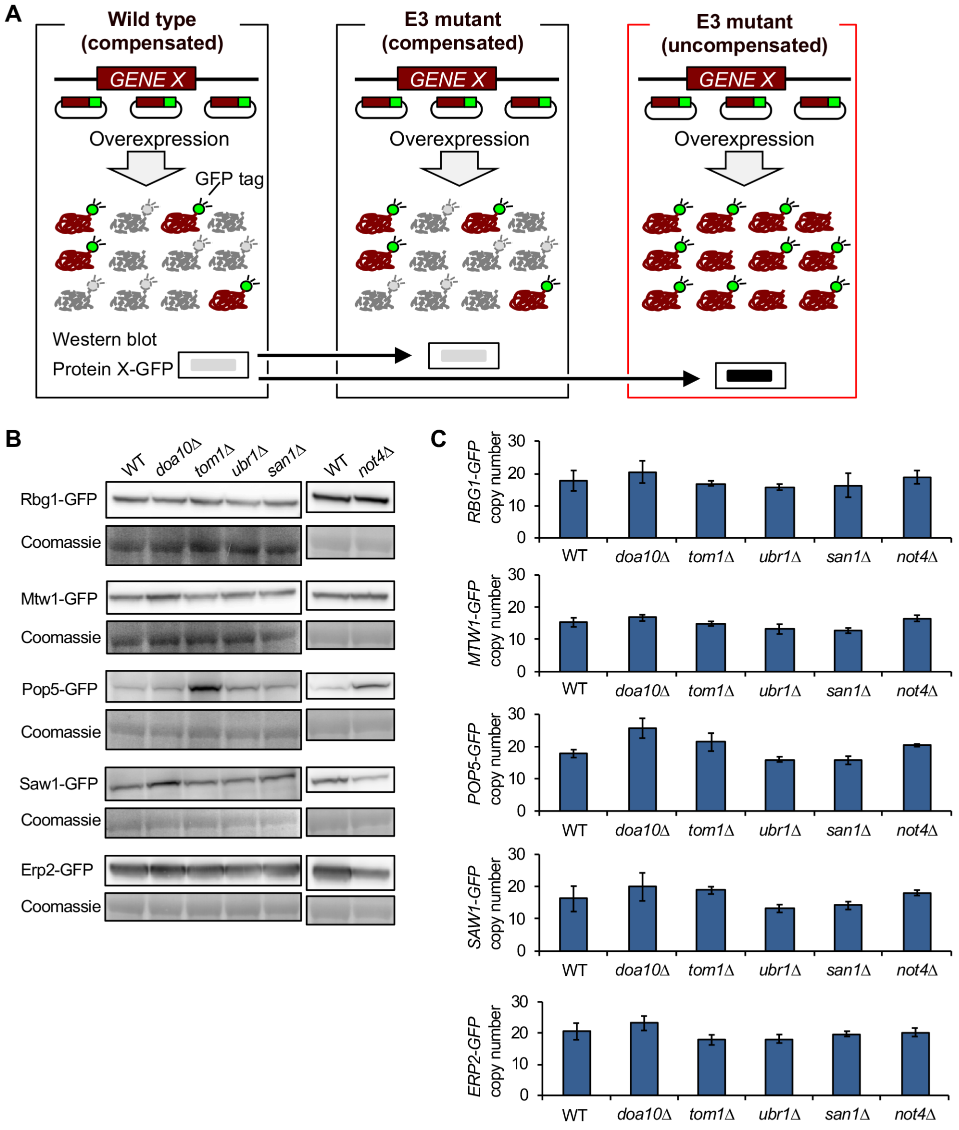


**S1 Fig. Identification of E3 ubiquitin ligases involved in dosage compensation.** (**A**) An experimental setup for the screen of E3 ubiquitin ligases involved in dosage compensation. The dosage-compensated proteins tagged with green fluorescent protein (GFP) were expressed from multicopy plasmid pTOW40836 containing the native regulatory sequences, including promoter and 5’ and 3’ untranslated regions. If the tested E3 ligase is not responsible for degradation of the target protein, the protein level is the same between WT and E3 mutant cells (left and middle panels). On the other hand, if the target protein is degraded through the tested E3 ligase, the protein level increases in the E3 mutant compared to WT cells (right and left panels). (**B**) Western blot of the GFP-tagged dosage-compensated proteins in E3 mutants using anti-GFP antibody. Coomassie staining of a 50-kDa protein, corresponding to enolase, is shown as a loading control. (**C**) Gene copy number during dosage compensation. Western blot detected the increased amount of Pop5 in *tom1*Δ and *not4*Δ and Saw1 in *doa10*Δ compared to those in WT cells, although the plasmid copy number was almost the same among the tested strains. Thus, Tom1 and Not4 and Doa10 were identified as E3 ubiquitin ligases involved in degradation of Pop5 and Saw1, respectively. Bar graph represents the copy numbers of pTOW40836 carrying each of the indicated genes in WT or E3 mutants. The average copy numbers ± s.d. were calculated from four technical replicates.

**Supplementary Figure 2**

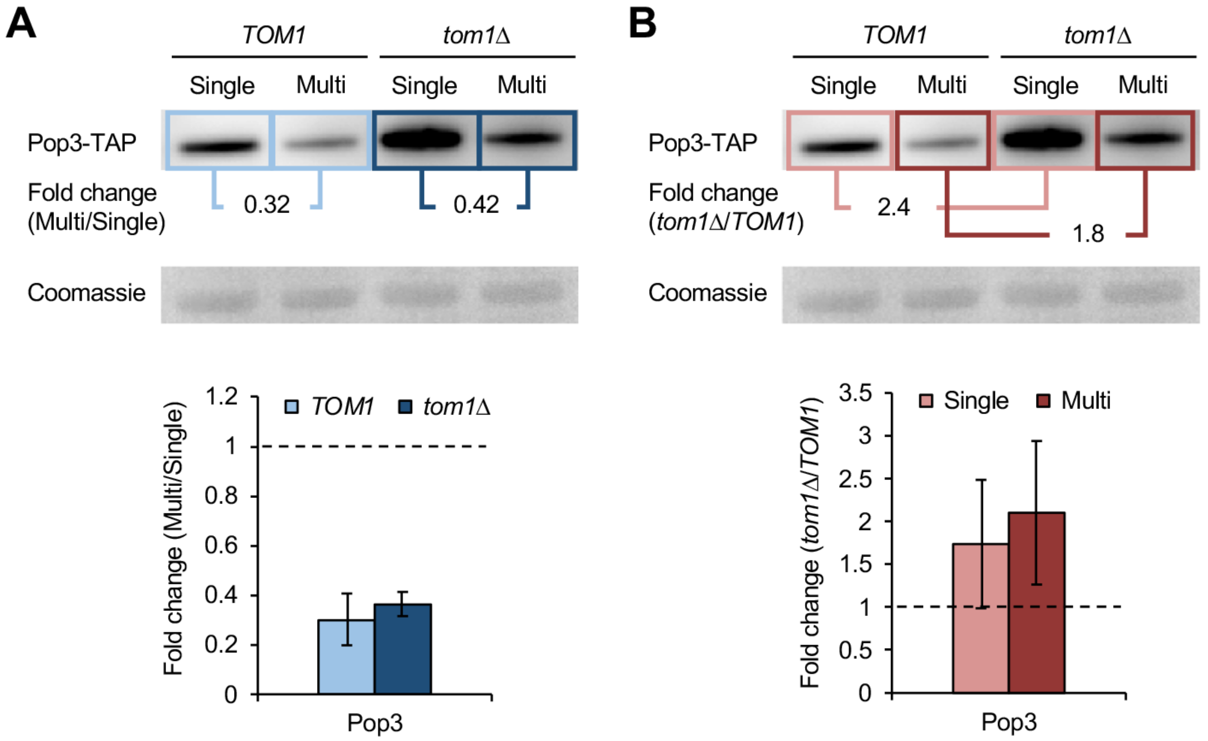


**S2 Fig. Quantification of Western blot data.**
(**A**) Comparison of protein levels between the Single and Multi conditions in WT or each mutant. Shown as an example is Pop3-TAP in WT and *tom1*Δ cells. The band intensity of Pop3-TAP in Multi was divided by that in Single in each strain. Coomassie staining of a 50-kDa protein, corresponding to enolase, is shown as a loading control. Data are from Fig 2A and 2B. (**B**) Comparison of protein levels between WT and each mutant in the Single or Multi conditions. The band intensity of Pop3-TAP in *tom1*Δ was divided by that in WT cells in each copy number condition.

**Supplementary Figure 3**

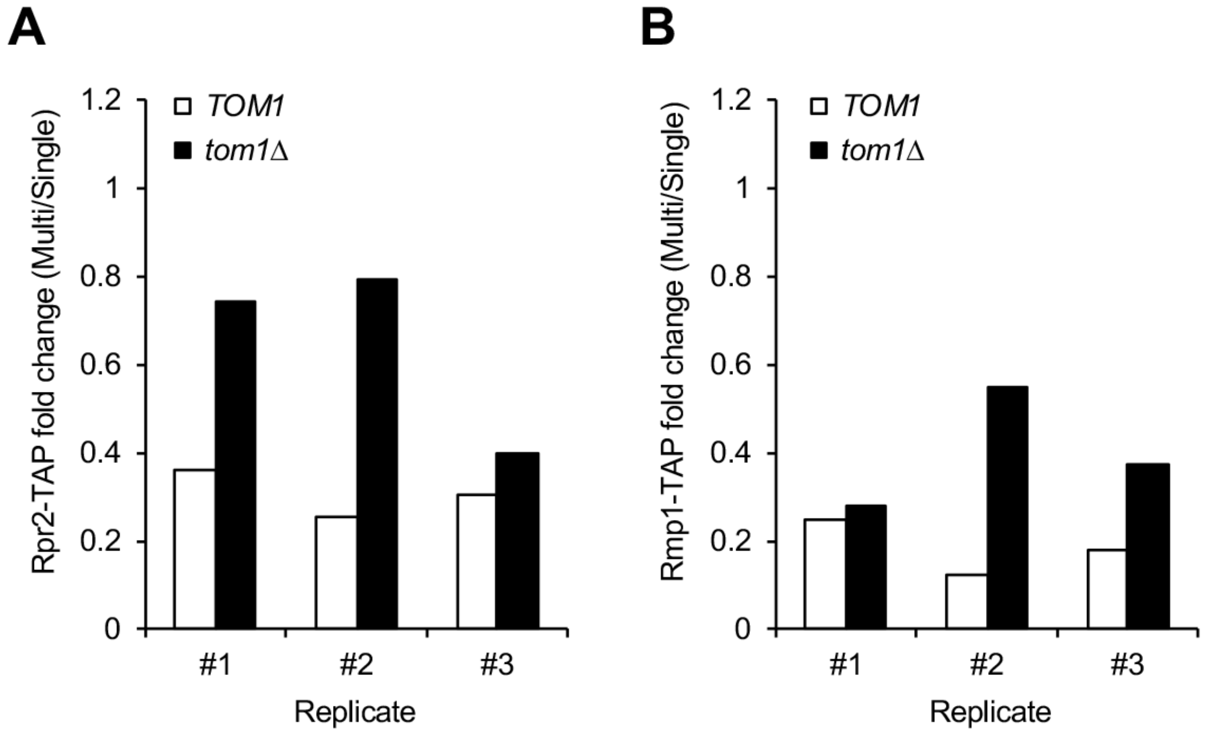


**S3 Fig. Rpr2 and Rmp1 tend to be less compensated in *tom1*Δ cells.**
(**A, B**) Lower compensation of Rrp2 (**A**) and Rmp1 (**B**) in *tom1*Δ cells was observed in three biological replicates. Quantification in each replicate is shown. Data are from Fig 2A and 2B.

**Supplementary Figure 4**

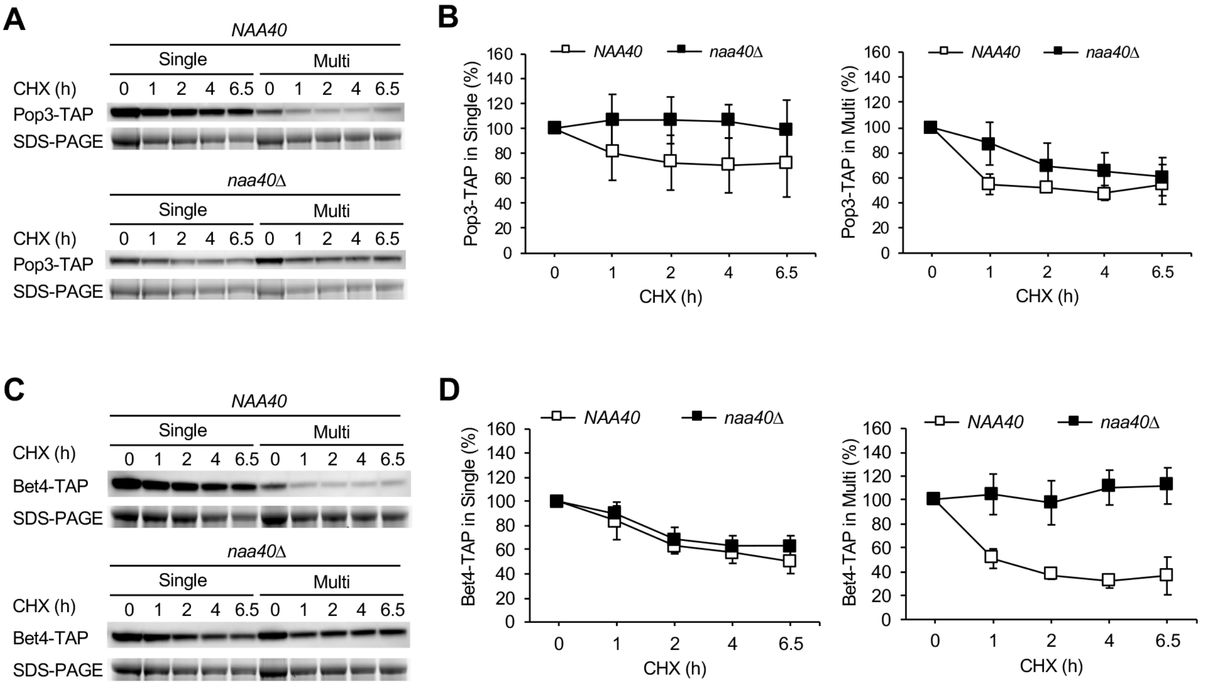


**S4 Fig. CHX chase experiments of Pop3-TAP and Bet4-TAP in *naa40*Δ cells.**
(**A, B**) CHX chase experiments of Pop3-TAP in WT and *naa40*Δ cells. Western blot with PAP and SDS-PAGE of a 50-kDa protein as a loading control (**A**). Quantification of Pop3-TAP levels in the Single (left) or Multi (right) conditions (**B**). The average protein level ± s.d. was calculated from three biological replicates. (**C, D**) Same as in (**A, B**), except that shown are Western blot and quantification of Bet4-TAP.

**Supplementary Figure 5**

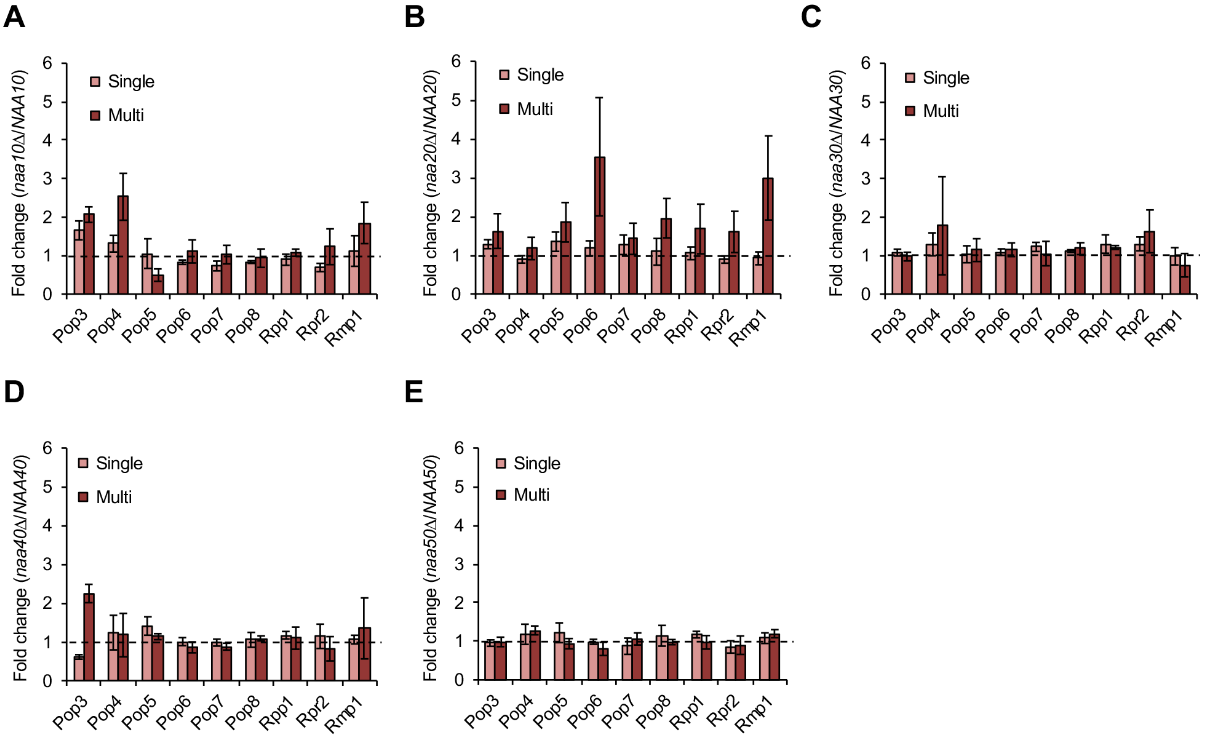


**S5 Fig. The effect of NATs on the endogenous protein level of the RNase P/MRP subunits.**
(**A–E**) Comparison of protein levels between WT and *naa10*Δ (**A**), *naa20*Δ (**B**), *naa30*Δ (**C**), *naa40*Δ (**D**), or *naa50*Δ (**E**) cells in the Single or Multi conditions. The average fold change ± s.d. was calculated from three biological replicates. Dashed line represents the same expression level between WT and mutant cells. Data are from Fig 4B–4K.

**Supplementary Figure 6**

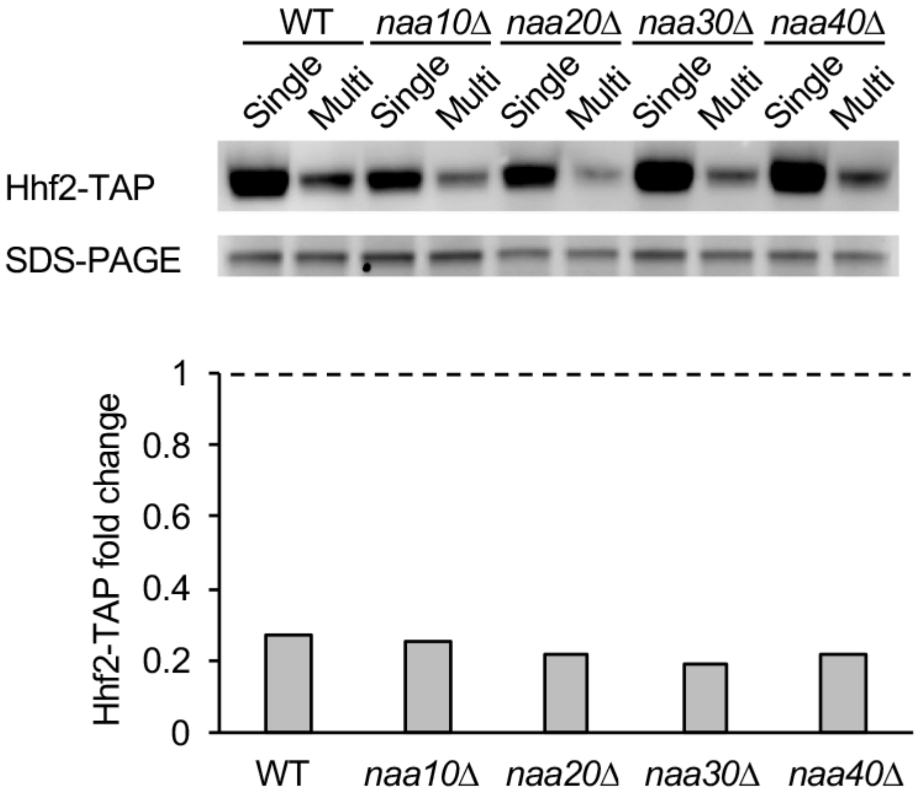


**S6 Fig. Dosage compensation of Hhf2 in NATs mutants.**
Western blot analysis of Hhf2-TAP in WT, *naa10*Δ, *naa20*Δ, *naa30*Δ, and *naa40*Δ cells. The TAP-tagged subunits were detected with PAP (top) and quantified (bottom). Dashed line represents the same expression level between the Single and Multi conditions.

**Supplementary Figure 7**

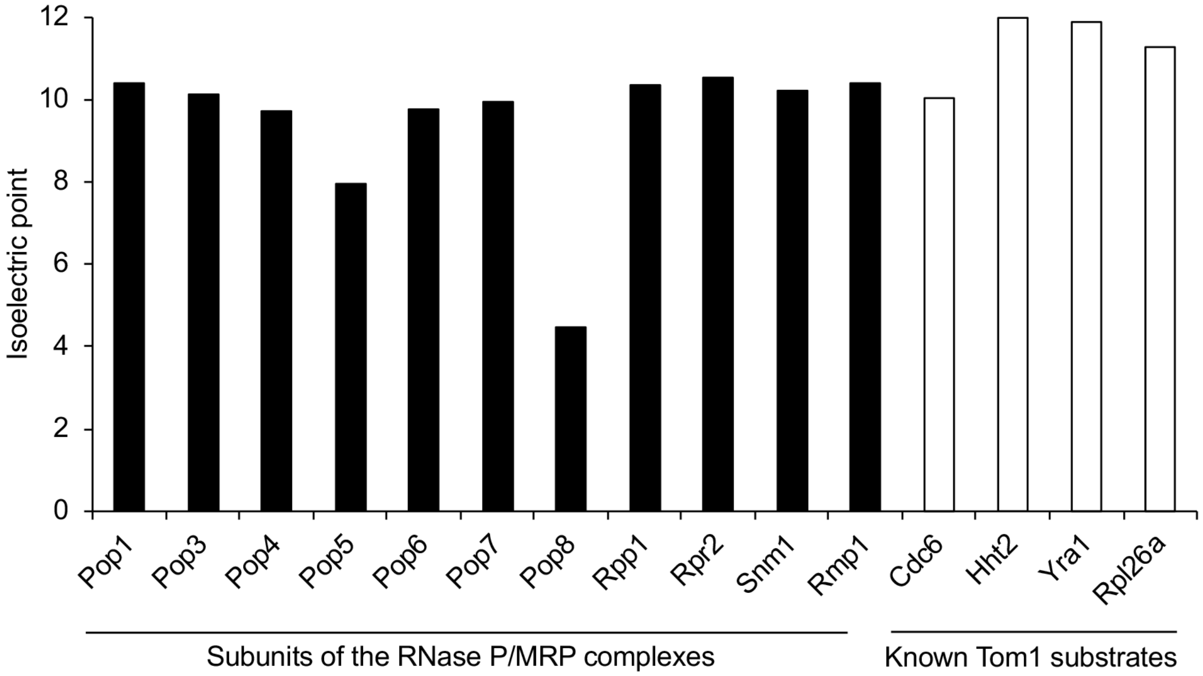


**S7 Fig. The isoelectric point of the RNase P/MRP subunits and Tom1 substrates.** The isoelectric point of Tom1 is 4.8, while that of the RNase P/MRP subunits, except Pop8, and known Tom1 substrates is around 10.
